# An ornithological survey of Fergusson Island, D’Entrecasteaux Archipelago, Papua New Guinea, reveals new island records and noteworthy natural history observations

**DOI:** 10.1101/2024.07.03.601923

**Authors:** Jordan Boersma, Jason Gregg, Doka Nason, Eli Malesa, Cosmo Le Breton, Serena Ketaloya, Bulisa Iova, John C. Mittermeier

## Abstract

Fergusson Island lies off the southeastern end of New Guinea and is the largest landmass in the D’Entrecasteaux and Trobriand Island Endemic Bird Area. We conducted audiovisual and camera trapping surveys in eastern Fergusson in September 2022 and recorded 97 bird species, documenting breeding and vocalizations for several of the EBA’s endemic taxa. Notably, we provide the first confirmed documentation of the “lost” *insularis* subspecies of Pheasant Pigeon *Otidiphaps nobilis* (often recognized as a distinct species, Black-naped Pheasant-pigeon), in 126 years, observations of eight bird species not previously recorded from Fergusson, and nesting of Goldie’s Bird-of-Paradise *Paradisea decora* and the endemic *crookshanki* taxon of Capped White-eye *Zosterops fuscicapilla* (or Oya Tabu White-eye). Our new distributional records were mostly either migratory species or species found in open habitats, highlighting the value of surveying across seasons and habitats. We summarise our results and provide comments on distributional records, breeding behaviour, and vocalizations.

---

Located off the south-eastern end of New Guinea, the D’Entrecasteaux Archipelago in Milne Bay Province, Papua New Guinea, forms part of the D’Entrecasteaux and Trobriand Islands Endemic Bird Area (BirdLife International 2024a, Stattersfield *et al*., 1998). The archipelago consists of three main islands, Goodenough, Fergusson, and Normanby, with the middle island in the chain, Fergusson, being the largest of the three (1,437 km^2^). High levels of endemism in this archipelago are the product of these being true oceanic islands that were never connected to the New Guinea mainland (except potentially part of Normanby, see Hill et al. 2023) and are separated by 18 km at their closest point from mainland New Guinea. The bird communities of these islands overlap substantially with mainland New Guinea (Mayr & Van Deusen 1956) but are less diverse due to the dispersal barrier presented by the open water channel between the mainland and the islands (Diamond 1972). The EBA is home to three endemic species: Long-billed Myzomela *Myzomela longirostris*, Curl-crested Manucode *Manucodia comrii*, and Goldie’s Bird-of-Paradise *Paradisea decora*, as well numerous endemic subspecies including two, the *insularis* subspecies of Pheasant Pigeon *Otidiphaps nobilis* and the *crookshanki* subspecies of Capped White-eye *Zosterops fuscicapilla*, which are sometimes treated as distinct, endemic species, Black-naped Pheasant-pigeon and Oya Tabu White-eye, respectively (Pratt & Beehler 2015, HBW/BirdLife Taxonomic Checklist v8.1).

Despite the ornithological significance of the D’Entrecasteaux and Trobriand Islands, research on the birds in this region has been limited and scientific knowledge of the EBA remains poor (BirdLife International 2024a). The first ornithological collections from the islands were in the late 1800s with notable collections being made by Andrew Goldie (for whom the endemic bird-of-paradise is named) in 1882 along with multiple visits to the islands by Albert S. Meek between 1894 and 1913 (see Frith & Beehler 1998 and Gregg et al. 2020 for a summaries of ornithological visits). The most recent ornithological study on Fergusson was a two-week survey of the south-western slope of the Oya Tabu Massif in August 2019 by JG, DN, and JB (Gregg et al. 2020). The fact that this study contributed five new records for the island, including four breeding species, provided evidence that even basic information such as the overall breeding bird diversity remains incomplete for islands in the D’Entrecasteaux Archipelago.

Even less is known about the natural history, vocal behaviour, and conservation threats for endemic taxa to the EBA. As a notable example, there is apparently no existing description of the nest of Goldie’s Bird-of-Paradise despite this being a charismatic and relatively common species on the island. Likewise, recordings and descriptions of the vocalizations of several endemic taxa, such as the *ochracea* subspecies of Yellow-billed Kingfisher *Syma torotoro, fortis* subspecies of Variable Shrikethrush *Colluricincla fortis*, and *finschii* subspecies of South Papuan Pitta *Erythropitta macklotii*, are limited. In addition to providing basic ecological information on species, these data are relevant for taxonomic evaluations and understanding potential conservation threats to birds in the region.

We surveyed areas in the eastern half of Fergusson Island, with a specific focus around the Oya Tabu Massif, between 5-30 September 2022. This work built on the 2019 survey by JG, DN, and JB and formed part of a larger project to assess the conservation status of the *insularis* subspecies of Pheasant Pigeon. Here we report the results from those field surveys and provide comments on our records of Pheasant Pigeon (see also Gregg et al., *in prep*), new distributional records for Fergusson Island, and observations of breeding behaviour and the vocalizations of endemic taxa.

## Methods

We visited eastern Fergusson Island from 5-30 September 2022. Our survey locations included three communities around the Oya Tabu Massif, Basima (S 9.4660, E 150.8331), Sion (S 9.4480, E 150.8095) and Bosalewa (S 9.4460, E 150.7205), and four communities in Fergusson’s eastern lowlands, Salamo (S 9.6662, E 150.7939), Galubwa (S 9.6125, E 150.7816), Upper Momoawa (S 9.5598, E 150.8148) and Bibio on Sebutuia Bay (S 9.5754, E 150.8635; see Figure 1). In addition to visiting habitats around these communities, we conducted surveys at three camps in primary forest ca. 1 day walk from the nearest community: on Mt Oya Tabu above Sion village (S 9.4554, E 150.7888, 1,110 m, 7-11 September); Kalatupe near Bosalewa (S 9.4535, E 150.7451, 525 m, 23-25 September), and on the Kwama River south of Bosalewa (S 9.5128, E 150.7584, 860 m, 26-28 September). These camps were on the northern, western and southern, respectively, slopes of the Oya Tabu Massif.

**Figure 1.**
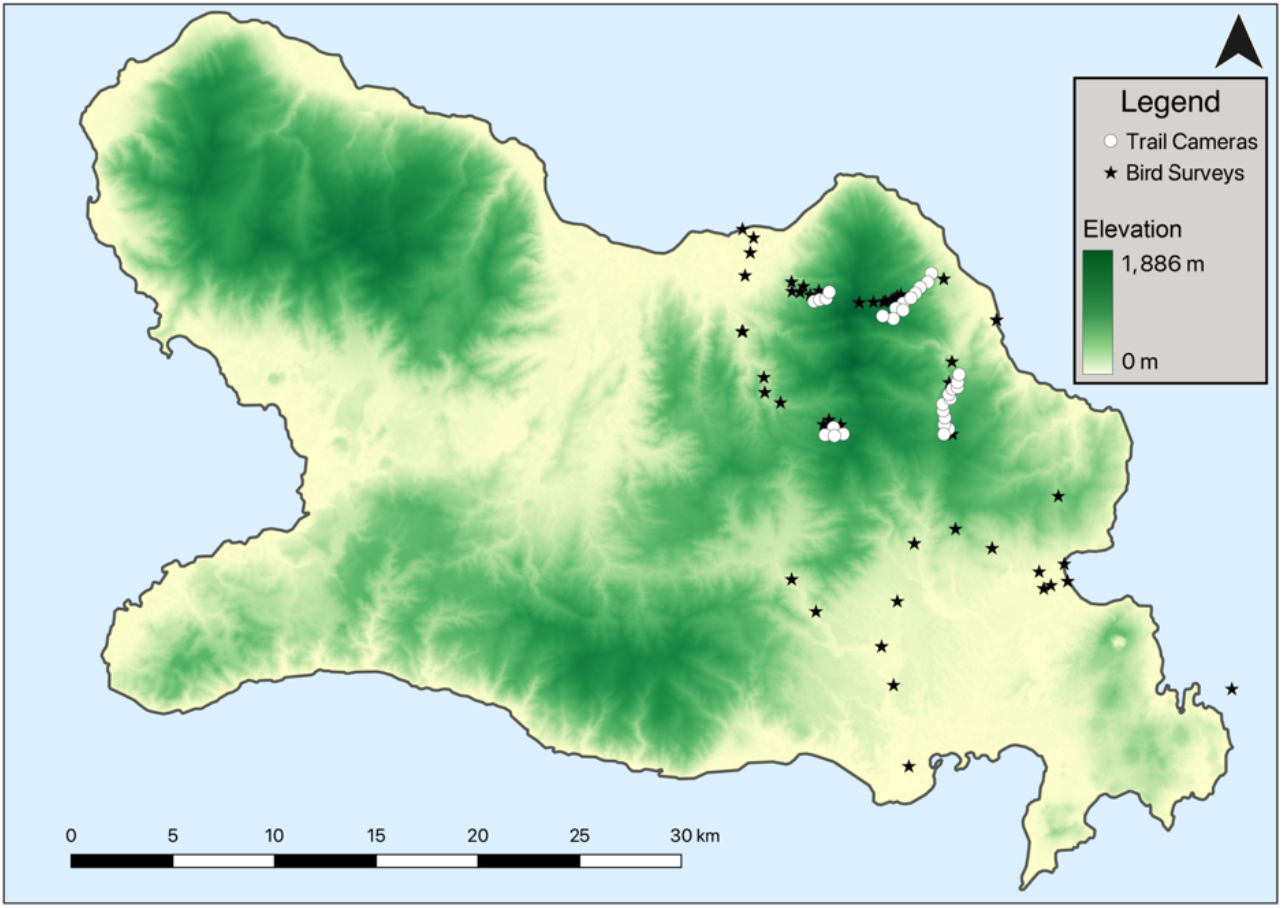
Map of survey and camera trap locations on Fergusson Island, Milne Bay Province, Papua New Guinea. Some camera locations are jittered from their actual location to reduce overlap with bird survey points.

Oya Tabu, also known as Mt Kilkerran, is the largest and highest mountain on Fergusson Island (max elevation 2,073 m) and forms part of a massif with several lower surrounding peaks such as Mt Olaba on its northern edge. Vegetation on the massif constitutes a mosaic of forest fragmented by subsistence agriculture around villages and logging, to an altitude of ca. 300 m (see Fig. 2 for habitat photographs). The forest is mostly undisturbed above 600 m, and montane cloud forest is characterised by epiphytes, epiphylls, and stands of bamboo at altitudes over 1,300 m.

**Figure 2.**
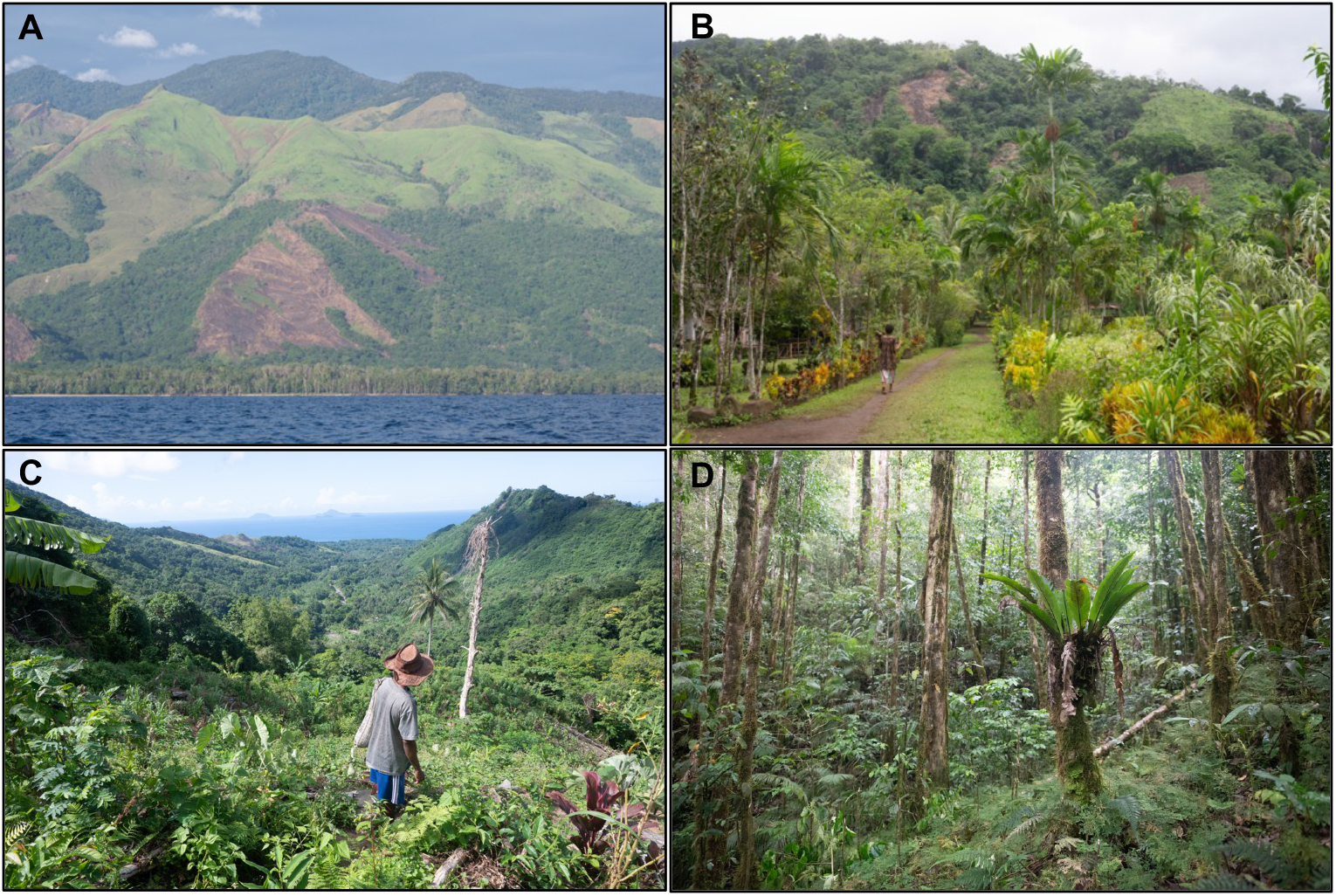
Representative images for habitats surveyed on Fergusson Island: **A**. Fergusson Island from sea level to higher mountains, showing gradient of human-altered and primary forest ecosystems. **B**. Human-modified habitat around villages containing betel nut palms and ornamental plants. **C**. A garden on the slopes of Mt. Oya Tabu containing subsistence crops including banana, tapioca, and taro. **D**. Upper montane habitat on Oya Tabu, showing understory of ferns and trees blanketed in moss and epiphytes. Photos by JCM.

To evaluate which species had previously been reported on Fergusson, we reviewed published records from the D’Entrecasteaux Archipelago (e.g., Beehler & Pratt 2016) and reviewed specimens in the American Museum of Natural History (New York, USA).

### Bird surveys

We conducted bird surveys on all 26 days we were present on Fergusson. At our forest camps, surveys began at dawn and continued until late morning with additional outings in the late afternoon and evening. At sites around villages, we did morning walks accompanied by local community members. Surveys were supplemented by opportunistic observations whenever possible. We recorded species using the eBird mobile app following either stationary (N = 26), traveling (N = 37), or incidental (N = 7) count methods (Sullivan et al. 2009). Locations and survey distances were recorded with the GAIA GPS application (www.gaiagps.com) or a Garmin 60CSX GPS. We used a Sennheiser ME66 microphone and Marantz Solid State Recorder (PMD661) to record vocalisations, and dslr cameras with telephoto lens to photograph species, when possible.

### Camera traps

We deployed eight camera traps along an elevational transect from 360 m to 1,360 m asl along the northern slope of the Oya Tabu Massif above Sion village between September 7-11, and twelve cameras along a similar elevational transect between 360 m and 970 m above Basima village from September 13-30. Cameras were placed a minimum of 200 m distance apart, alternating between Browning Strikeforce Pro BTC-5DCL (Birmingham, AL, USA) and Reconyx Hyperfire H2X (Holmen, WI, USA) models along the transect. From September 26-28 we deployed an additional eight cameras in primary forest between 550 and 980 m around our Kwama River camp. In all areas, camera traps were set in sites with little or no human disturbance, prioritising game trails and places with recently fallen fruit when possible. Cameras were mounted vertically on PVC poles, placed ca. 10 cm above the ground, and set to high sensitivity and rapidfire photo bursts. We also opportunistically deployed cameras near nests for up to 24 hours to confirm species and record nest behaviour. For full details on camera trap methods, see Gregg et al. (*in prep*).

## Results

We recorded 97 bird species during our surveys on Fergusson Island and estimated abundance for each based on the number of detections during audiovisual surveys and camera trap hours in suitable habitat (see Table 1). Audiovisual surveys detected 95 species compared to eight species photographed by camera traps (Figure 3). Pheasant Pigeon and Red-necked Crake *Rallina tricolor* were the only species that were recorded with camera traps but not observed during the audiovisual surveys. Orange-footed Megapode *Megapodius reinwardt* and South Papuan Pitta accounted for the majority of bird detections by the camera traps overall (36% and 28% of bird detections, respectively).

**Table 1.**
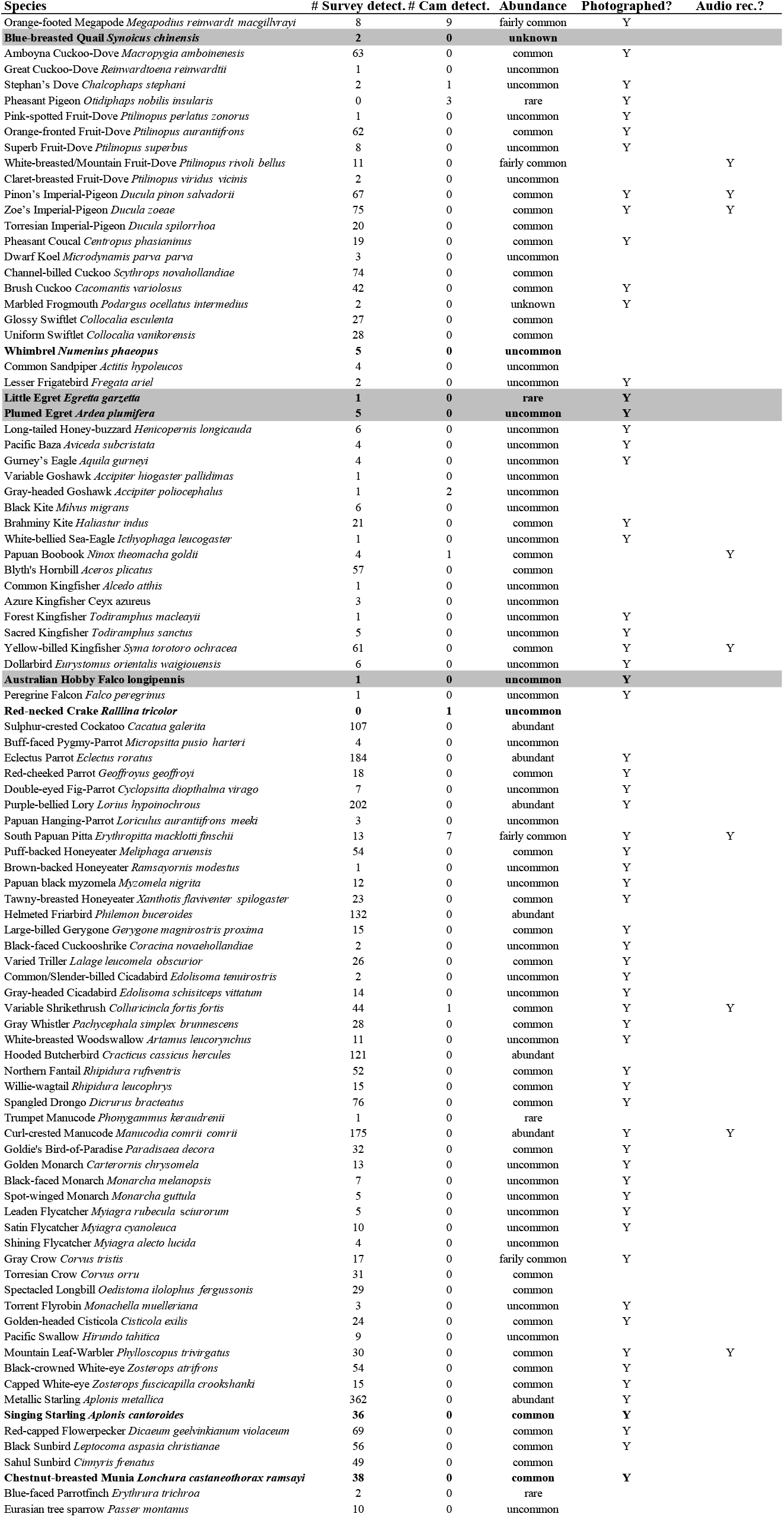
Species recorded during field surveys between 5-30 September 2022 in Fergusson Island, Milne Bay Province, Papua New Guinea. Bolded species reflect new records for Fergusson Island and highlighted rows depict new records for the D’Entrecasteaux Archipelago; where appropriate, local subspecies is listed. We estimated number of individuals for each species during on-foot surveys, totalled the number of detections from trail cameras, and provided general descriptions of relative abundance based on the number of detections relative to effort in suitable habitat for each species. All photographs and audio recordings can be found on Macaulay Library (https://www.macaulaylibrary.org/).

**Figure 3.**
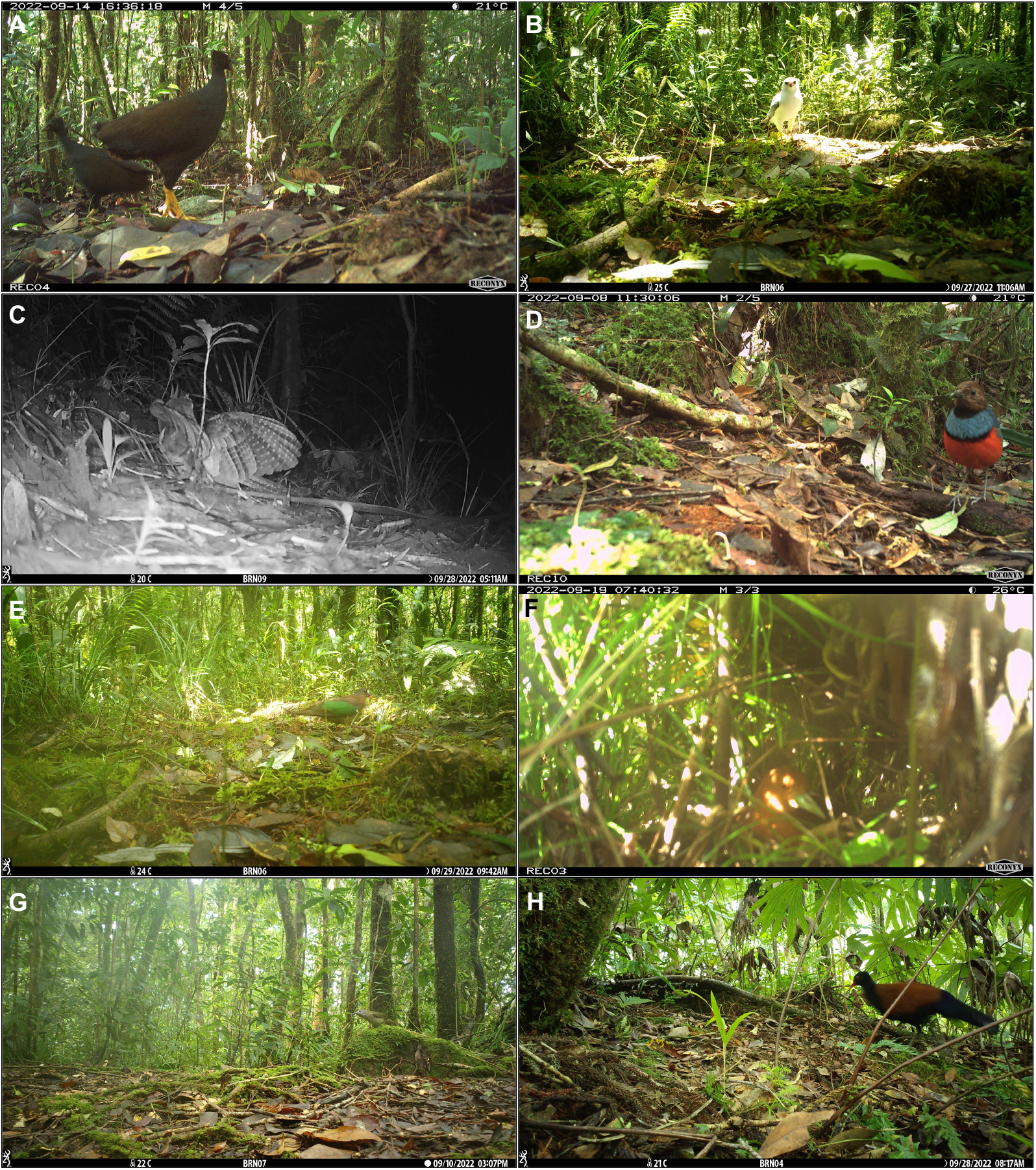
Trail camera images for: **A**. Orange-footed Megapode *Megapodia reinwardt macgillivrayi*, **B**. Gray-headed Goshawk *Accipiter poliocephalus*, **C**. Papuan Boobook *Ninox theomacha goldii*, **D**. South Papuan Pitta *Erythropitta macklotti finschii*, **E**. Stephan’s Dove *Calcophaps stephani*, **F**. Red-necked Crake *Rallina tricolor*, **G**. Variable Shrikethrush *Colluricincla fortis fortis*, and **H**. Pheasant Pigeon *Otidiphaps nobilis insularis*.

The majority (74%) of the 97 species we recorded are documented by media vouchers (photographs or sound recordings) all of which are archived and publicly available in Macaulay Library (https://www.macaulaylibrary.org/; see Table 1). Comments on the *insularis* subspecies of Pheasant Pigeon, new island records for Fergusson, noteworthy breeding observations, and vocalizations of endemic taxa are provided below. Taxonomy and nomenclature of the species listed follows Clements/eBird v2023 (Clements *et al*., 2023), which matches the taxonomy used by Macaulay Library where our supporting media is archived.

### Pheasant Pigeon *Otidiphaps nobilis insularis*

The *insularis* subspecies of Pheasant Pigeon is known only from Fergusson Island and is treated as a distinct, and critically endangered, species, Black-naped Pheasant-pigeon, by the IUCN Red List and HBW/BirdLife Taxonomic Checklist (BirdLife International 2024b). Prior to our fieldwork, the taxon was known from only three specimens: two individuals collected by Goldie in 1882, which formed the type series, and a single specimen collected by Meek in 1896 (Kirwan et al. 2023). With no confirmed documentation of this taxon in 126 years, Black-naped Pheasant-pigeon was identified as a “lost” bird species by the Search for Lost Birds (Rutt & Mittermeier 2023) and provided the impetus for much of our survey and camera trap efforts (see Gregg et al., *in prep*).

We detected Pheasant Pigeon on camera traps on three occasions in September 2022: a single individual photographed by camera set on the north slope of Oya Tabu above Basima village(746 m) on 22 September, and presumably one individual captured on a camera near our Kwama River camp on 27 and 28 September (971 m, Fig. 3H). The fact that we only detected this species twice despite a significant search effort indicates that the population on Fergusson is both rare and extremely elusive and that, using a precautionary approach, its Red List status of critically endangered is warranted. The two locations where the bird was found were both in relatively undisturbed primary hill forest where there had not been any logging activity. Conversations with local residents, however, suggested that at least one of these areas could be designated for logging in the near future.

### New Distributional Records for Fergusson Island

We recorded eight species that do not have previously published records from Fergusson Island, including five of which that do not have published records for anywhere in the D’Entrecasteaux Archipelago (Figure 4). These included two austral migrants to New Guinea (Little Egret *Egretta garzetta* and Australian Hobby *Falco longipennis*), one boreal migrant (Whimbrel *Numenius phaeopus*), four species that are likely resident in open and human-modified habitats (Blue-breasted Quail *Synoicus chinensis*, Plumed Egret *Ardea plumifera*, Singing Starling *Aplonis cantoroides*, Chestnut-breasted Munia *Lonchura castaneothorax*) and one breeding forest species (Red-necked Crake *Rallina tricolor*).

**Figure 4.**
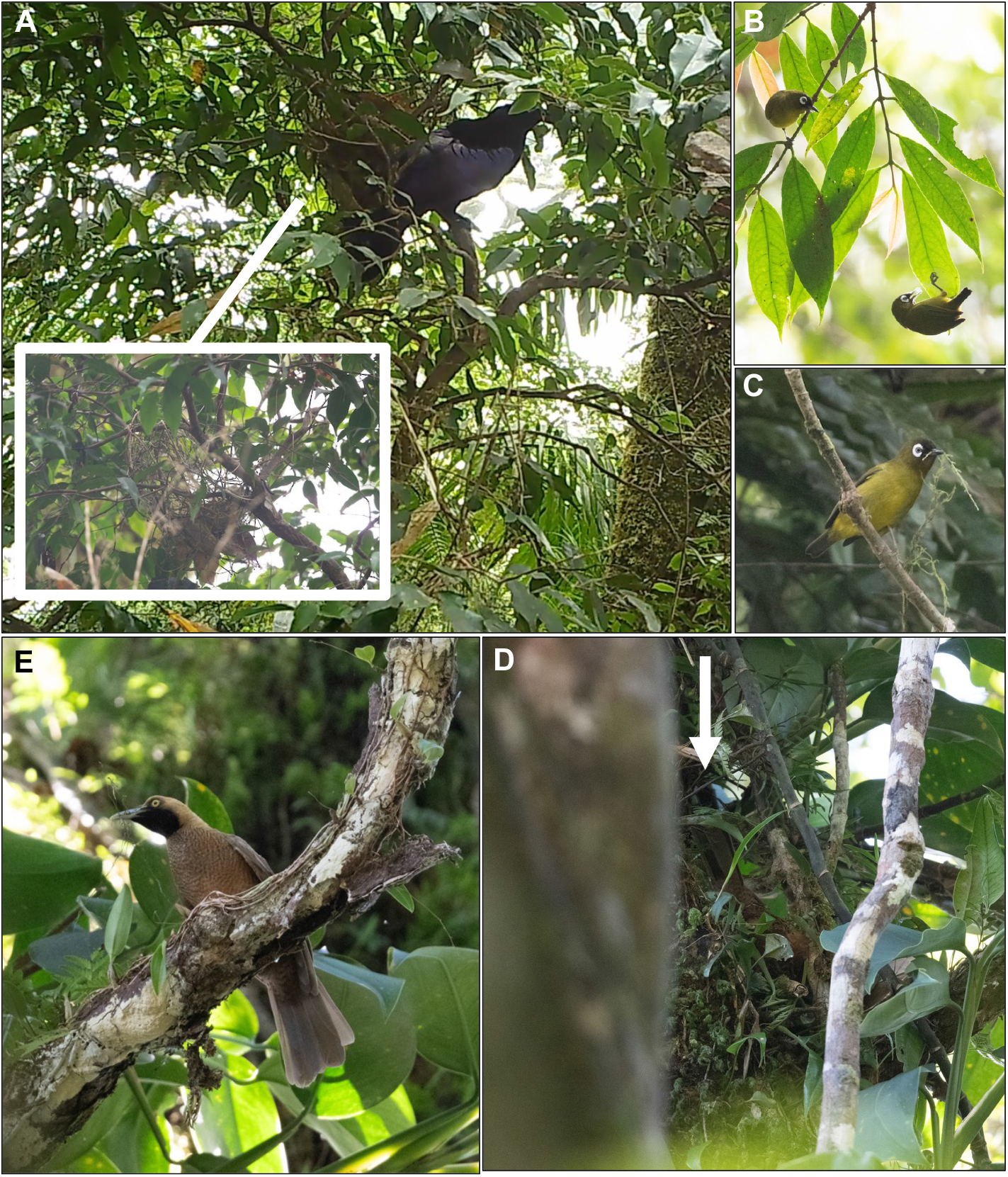
Nest-building behaviour and nests of select species: **A**. Curl-crested Manucode *Manucodia comrii comrii* visiting nest and inset image of nest in building stage, **B-C**. *crookshanki* subspecies of Capped White-eye *Zosterops fuscicapilla* collecting nesting material, **D**. Goldie’s Bird-of-Paradise *Paradisea decora* female carrying nesting material, and **E**. Goldie’s female on a nest in an epiphyte (arrow points to tail). Photos by DN (A) and JCM (B-E)

#### Blue-breasted Quail *Synoicus chinensis*

On 22 September, JG flushed two quail from open grassland at sea level near Idubada station, Bosalewa community, and made an audio recording of the birds calling at close range (ML610283282, see Appendix 1 for spectrogram). Based on the recorded vocalizations, we identified the birds as Blue-breasted Quail. Blue-breasted Quail occur in lowlands habitats in most of New Guinea and in the Bismarck Archipelago, but had no previously published records from the D’Entrecasteaux Islands.

#### Red-necked Crake *Rallina tricolor*

On 18 September, a local resident in Galubwa directed JB and DN to a bird nest he had found while clearing a garden and a trail camera we placed near the nest subsequently documented an adult Red-necked Crake coming to incubate (Figure 3F). Red-necked Crake is a widespread species on the islands around New Guinea with a previous record from Normanby Island (Beehler & Pratt 2016), making the presence of this species as a breeder on Fergusson unsurprising. Notably, this was the only breeding forest species that we detected that had not previously been reported from the island.

#### Whimbrel *Numenius phaeopus*

We observed small groups of Whimbrel on September 14 in Sebutuia Bay. This species is speculated to be the most common shorebird throughout New Guinea (Bishop 2006), and has previously been detected on Goodenough Is. (Bell 1970), so its occurrence on Fergusson is unsurprising despite the lack of published records. The individuals we observed are likely migrants on their way to mainland NG or Australia.

#### Little Egret *Egretta garzetta and* Plumed Egret *Ardea plumifera*

We observed a single individual on Bwaghalu beach near Bosalewa village on 22 September. The timing of this record during the austral winter fits the known pattern of non-breeding dispersal from Australia. Plumed Egret breeds throughout the New Guinea region but has apparently not been documented from Fergusson before. We observed Plumed Egrets on multiple occasions between 17-21 September in open areas near Salamo, Galubwa, and Upper Momoawa. Neither Little nor Plumed Egret had published records from the D’Entrecasteaux Archipelago prior to our 2023 trip. Little Egret is a non-breeding visitor to wetlands in New Guinea from Australia. The lack of previous records for both species almost certainly results from these widespread species being overlooked or of little interest to previous ornithologists. Their presence on Fergusson suggests they likely occur throughout the archipelago.

#### Australian Hobby *Falco longipennis*

We observed a single Australian Hobby in open grassland habitat near Bosalewa village on September 22 (Figure 2E). This is the first confirmed record of this migratory species for the D’Entrecasteaux Archipelago. Similar to Little Egret, this species occurs in New Guinea as a non-breeding visitor from Australia in the austral winter. The lack of previous records from D’Entrecasteaux may be due in part to earlier surveys either overlooking migratory species or occurring in seasons when they were not present on the island.

#### Singing Starling *Aplonis cantoroides*

We found Singing Starling to be a common and conspicuous species around Salamo, the largest human settlement on Fergusson, and regularly observed flocks of up to ca. 20 individuals as well as recently fledged juvenile birds. This species is known to occur on neighbouring Normanby Is., but had not previously been documented elsewhere in the archipelago. We did not observe Singing Starling in other parts of east Fergusson outside of Salamo. Its congener, Metallic Starling *Aplonis metallica*, was commonly detected at other locations. Singing Starlings often occurs around human settlements and agricultural areas in the New Guinea islands, and while the species could have been overlooked by previous ornithologists, it may also be a new arrival to Fergusson that colonised the island following human-induced habitat change around Salamo.

#### Chestnut-breasted Munia *Lonchura castaneothorax*

We observed Chestnut-breasted Munias in open areas and secondary growth around Salamo on 16-17 September and in grasslands near Bosalewa on 22 September. Despite appearing to be a common in suitable habitat near villages, Chestnut-breasted Munia has not previously been documented on Fergusson. Similar to Singing Starling, it could be a recent colonist to the island or a species that has been overlooked given the limited information on birds in open habitats.

### Nesting Behaviour

We documented nests for three taxa endemic to the D’Entrecasteaux and Trobriand Islands EBA during our fieldwork.

#### Curl-crested Manucode *Manucodia comrii comrii*

Endemic to the D’Entrecasteaux and Trobriand Islands EBA, Curl-crested Manucodes were common on Fergusson from sea level to the highest elevations we visited on Oya Tabu. Nesting information for this species is summarized in Frith and Beehler (1998) and in Frith and Frith (2020). On 8 September we observed a Curl-crested Manucode constructing a nest in primary montane forest at ca. 1,250 m elevation near our camp on Mt Oya Tabu. The nest was located near the top of a ca. 8 m tall sapling and constructed of sticks and vines with a lining of dead leaves (see Fig. 4a).

#### Goldie’s Bird-of-Paradise *Paradisaea decora*

Unlike Curl-crested Manucode, published information on the nesting behaviour of the endemic Goldie’s Bird-of-Paradise is limited. Frith and Beehler (1998) note that males in breeding condition have been collected in September and that information on the nest itself is limited to second-hand accounts from local people who reported that “birds make a hole in a bird’s nest fern *Asplenium* sp.” The authors note that “this unlikely nest site for a *Paradisaea* sp. requires confirmation” (Frith & Beehler 1998, p. 474). We detected Goldie’s Birds-of-Paradise daily in the appropriate habitat including multiple displaying males in the lowlands near Bibio on Sebutuia Bay. On 9 September, JG and JCM observed a female collecting ca. 30-50 cm long strands of plant fibre and using it to construct a nest (see Fig. 4D). The nest was in primary hill forest at 986 m near our camp on Mt Oya Tabu and was located ca. 20 m above the ground on top of a clump of epiphytes. In addition to collecting material, the female was also observed sitting on the nest (See Fig. 4E) suggesting that she may have already been incubating. Due to its location, it was not possible to see the construction of the nest itself. The height and placement of this nest is similar to published information for other *Paradisaea* sp. (Frith and Beehler 1998) and the timing of bird nesting in September fits the, albeit limited, published information about the breeding phenology of this species. Female Raggiana Bird-of-Paradises *P. raggiana* have been observed adding vines and plant fibers to nests during incubation (Frith & Beehler 1998), and this may have been a similar observation. Goldie’s Bird-of-Paradise is considered vulnerable by the IUCN Red List due to a declining population limited to the remaining primary rainforest on Fergusson and Normanby Islands (Birdlife International 2024b). While we observed this species regularly in suitable forest, it was absent from areas directly around villages and areas with a heavy human presence.

#### Capped White-eye *Zosterops fuscicapilla crookshanki*

Capped White-eye occurs in highland habitats across mainland New Guinea, with the taxon *crookshanki* the only population of species complex occurring off the New Guinea mainland. Endemic to the highlands of Goodenough and Fergusson, *crookshanki* is often treated as a distinct species, Oya Tabu White-eye (e.g., Pratt & Beehler 2015), and little has been published about the ecology of this endemic taxon. Goodenough and Fergusson populations differ phenotypically, with the latter more closely resembling mainland Capped White-eye (T. Pratt, *pers. comms*.). We observed Capped White-eyes on several occasions between 1,200-1,800 m on the eastern flanks of Oya Tabu from September 8-10. On September 8 and 9, we observed pairs collecting nest material, and in one case building a nest ca.15 m above the forest floor in a ca. 25 m tall tree. In both observations we watched one individual collecting spider webs or moss while another individual mate-guarded (see Fig. 4B-C).

#### Vocalisations of Endemic Taxa

We recorded vocalisations for 12 species on Fergusson (Table 1). Among these are three taxa endemic to the D’Entrecasteaux Archipelago whose vocalizations have limited documentation. Since vocalizations are often helpful for determining species limits, we provide comments on these recordings and compare them to the vocalizations of mainland taxa.

#### Yellow-billed Kingfisher *Syma torotoro ochracea*

We found the endemic *ochracea* subspecies of Yellow-billed Kingfisher to be common at most of our survey locations from forest habitats at sea level near Salamo, Bibio, and Bosalewa to the highest elevations that we visited on Oya Tabu (ca. 1,800 m). This taxon been suggested as a potential species split from mainland populations on the basis of differences in morphology, call, and genetic divergence (Linck et al., 2020, Berryman et al., in prep). We recorded the calls of three individuals on 9 September (ML548009611 and ML548010441) and 15 September (ML513164643), nearly doubling known audio recordings of this taxon. Both the mainland and D’Entrecasteaux taxa exhibit a short and long version of their main call (see Fig. 5): the mainland taxon’s call is a fast trill, with pitch rising and falling once in the short and twice in the long version. The D’Entrecasteaux taxon’s is a considerably slower trill, with a slight vibrato in each note that steadily decreases in pitch and accelerates in the final notes. Long and short versions of this call differ only in that the longer version consists of a long repetition of one of the final notes in the call sequence.

**Figure 5.**
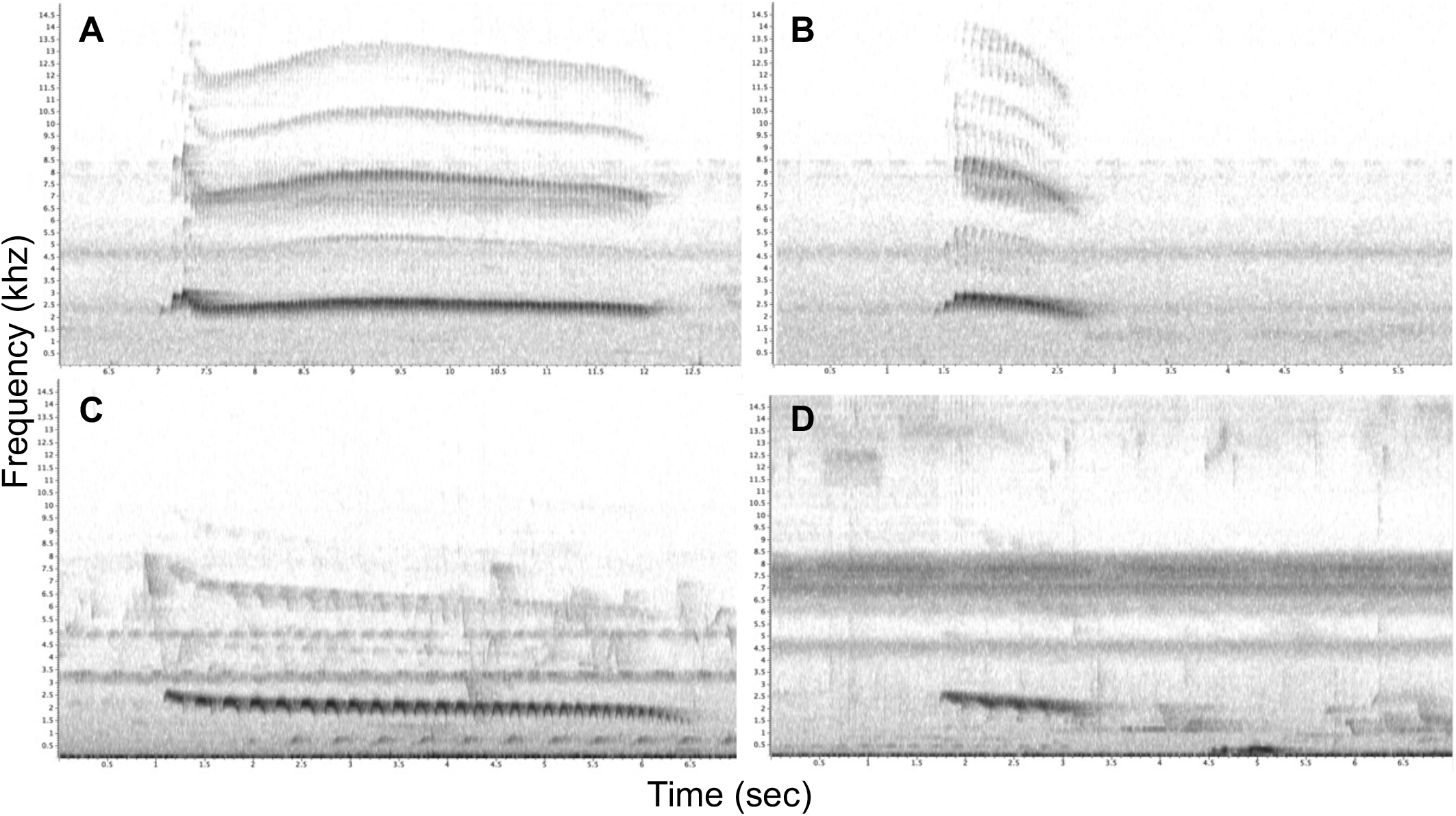
Spectrograms for Yellow-billed Kingfisher *Syma torotoro* vocalizations: **A**. long version of mainland *meeki* taxon (XC24796), **B**. short version of *meeki* taxon (XC38235), **C**. long version of endemic *ochracea* subspecies from our survey (ML548009611), **D**. short version of endemic *ochracea* subspecies from our survey (ML613164643).

#### South Papuan Pitta *Erythropitta macklotti finschii*

Recordings made by JB on 8 September (ML 547944611 and ML547945741) do not indicate substantial differences between the D’Entrecasteaux subspecies and the mainland taxa. Each call begins with a long, slowly upslurred tremulous note followed by a shorter, tremulous note that decreases slightly in pitch at the end.

#### Variable Shrikethrush *Colluricincla fortis fortis*

A recording made by JB on 9 September (ML 548004101) is the first recording of the *fortis* D’Entrecasteaux taxon and differs considerably from the few known recordings of the mainland *despecta* taxon, the only one of the four *C. fortis* taxa with published recordings. The *despecta* song differs across recordings, but always starts with at least one monotone introductory note, followed often by a downslurred or upslurred whistle (see Fig. 6). The *fortis* song typically consists of a sequence of three distinct downslurred notes, the first two being of the same approximate pitch and the last a higher pitch. Often the song alternates between the three-note sequence being of overall higher pitch followed by three notes of lower pitch. On occasion raspy notes of rising and falling pitch precede the standard song sequence.

**Figure 6.**
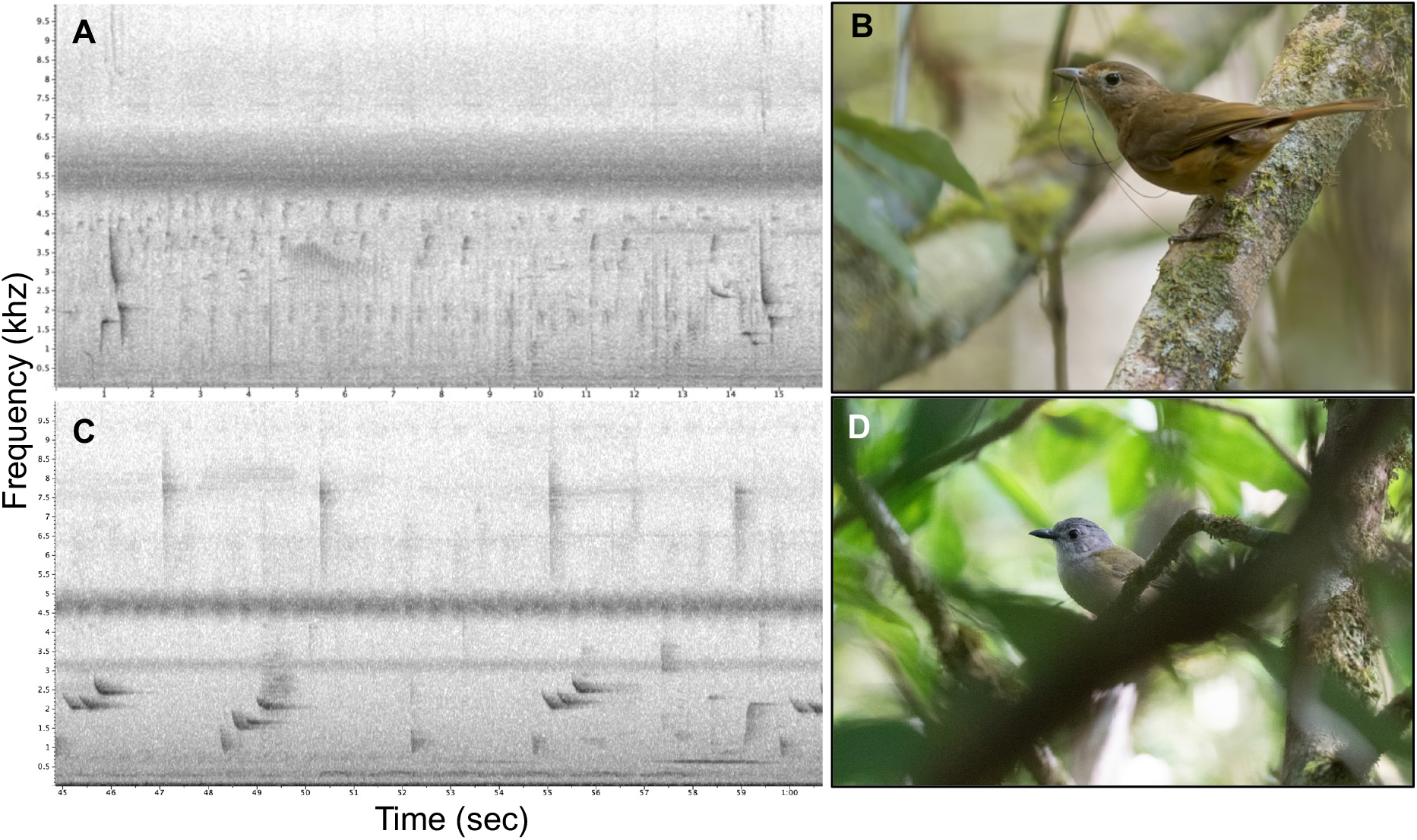
Comparison of spectrograms of Variable Shrikethrush *Colluricincla fortis* taxa: **A**. the typical song (XC279259) and **B**. adult plumage of mainland *despecta* taxon (ML575874601), and **C**. representative song (ML548004101) and **D**. adult plumage of the endemic *fortis* taxon from our surveys (ML533344161). Photos by Dubi Shapiro (B) and JCM (D).

## Discussion

We found published records for 121 species on Fergusson Island and 144 total species in the D’Entrecasteaux Archipelago (Appendix 2). Our field surveys added a further eight species that had no previously published records from Fergusson, increasing the total species recorded on the island from 121 to 129 (7% increase). Four of these species had no records from the D’Entrecasteaux Archipelago, increasing the total for the archipelago from 140 to 144 species (3% increase). Combined with the study by Gregg et al. (2020), recent surveys on Fergusson have increased the documented avian diversity of Fergusson by 11% and for the D’Entrecasteaux by 4%.

Of the eight new island records we found, three are migratory species (Australian Hobby, Little Egret, and Whimbrel), four are species found in savanna, secondary growth near human habitation, and gardens adjacent to villages (Blue-breasted Quail, Plumed Egret, Singing Starling, and Chestnut-breasted Munia), and one is an easily-overlooked species found in lowland forest that we documented with a camera trap (Red-necked Crake). Together these results highlight the value of surveying across seasons and habitats, as well as the benefit of using multiple survey methods, such as on-foot surveys, camera traps, and landowner engagement. Three of the species we recorded (Plumed Egret, Singing Starling, and Chestnut-breasted Munia), occurred only in the vicinity of villages and may constitute recent arrivals to Fergusson following human-led habitat change on the island. The recent record of Eurasian Tree Sparrow *Passer montanus* by Gregg et al. (2020) would also fit that pattern.

Camera traps allowed us to confirm the presence of two inconspicuous terrestrial species that had either not been recorded on the island before or had not been documented for more than a century: Red-necked Crake and Pheasant Pigeon. This demonstrates that while camera traps may not add significantly to the overall numbers of species detected during ornithological surveys, they can increase the detection of inconspicuous terrestrial birds, even when deployed for relatively brief periods of time. In our case, both species we documented with camera traps but not during audiovisual surveys were also reported by local people who described the birds and suggested locations for setting the cameras.

We regularly observed two of the three species endemic to the D’Entrecasteaux and Trobriand Islands EBA during our surveys: Curl-crested Manucode and Goldie’s Bird-of-Paradise, but failed to find the third endemic species, Long-billed Myzomela. This poorly-known species was originally described as endemic to montane forest on Goodenough Island and was first documented on Fergusson in 2019 (Gregg et al. 2020). While it appears to be uncommon on Fergusson, the fact that we failed to detect it during our survey period is likely due to the limited time we spent in forest above 1,000 m (four days total, including one with prohibitively bad weather conditions).

Birds on Fergusson and in the D’Entrecasteaux Archipelago face risks from climate change and habitat destruction. Two of the regionally endemic taxa (the *crookshanki* subspecies of Capped White-eye and Long-billed Myzomela) are restricted to the highest elevations and could face reduced suitable habitat as global climate continues to warm (Freeman et al. 2018). Logging has already degraded much of the lowland and hill rainforest and conversations with local landowners indicated that new operations are now targeting higher elevations with no prior disturbance. The most threatened bird on Fergusson is almost certainly the *insularis* subspecies of Pheasant Pigeon, which despite a significant effort we only detected on two occasions (see Gregg et al., *in prep*, for more details). This species is likely at risk from habitat degradation due to logging. Partnerships with local people were vital to the success of our fieldwork, as they guided us during surveys and shared their immense knowledge of local fauna. These local natural historians are ideal partners for developing conservation priorities before species decline or disappear.

## Acknowledgements

We thank the many landowners and local residents across East Fergusson that shared their unparalleled local natural history knowledge with us and granted access to their land for surveys. Our work was possible thanks to approval from the Papua New Guinea National Research Institute and the Milne Bay Provincial government. The project was funded by the RIDGES foundation with support from American Bird Conservancy. We are also grateful to D. Mitchell of Eco Custodian Advocates, T. Pratt, and G. Dutson for sharing their expertise on D’Entrecasteaux birds and logistics. T. Brooks helped with describing and visualizing vocalizations.

