## Appendix 1 for "An ornithological survey of Fergusson Island, D’Entrecasteaux Archipelago, Papua New Guinea, reveals new island records and noteworthy natural history observations"

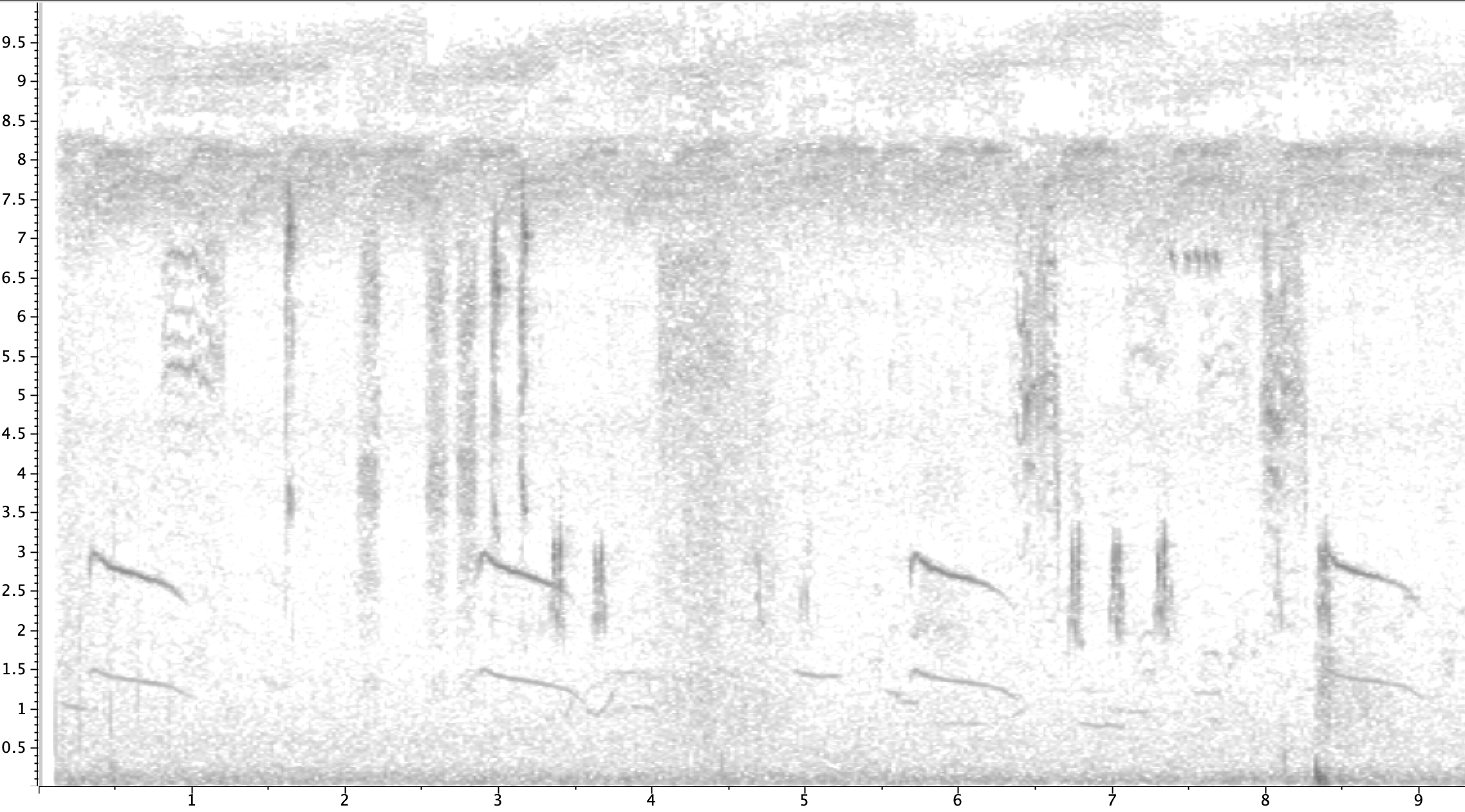

Frequency (khz)

Time (sec)

**Appendix 1.** Spectrogram for JG’s recording of Blue-breasted Quail Synoicus chinensis from Bosalewa village on 22 September, 2022 (ML610283282). This is the first confirmed record of this species on Fergusson Island.
