## Appendix 2 for "An ornithological survey of Fergusson Island, D’Entrecasteaux Archipelago, Papua New Guinea, reveals new island records and noteworthy natural history observations"

**Appendix 2.** Species list for the D’Entrecasteaux Archipelago, with presence/absence data indicated for each major island. Bolded species reflect new records for Fergusson Island and highlighted rows depict new records for the D’Entrecasteaux Archipelago from this expedition. Local subspecies is indicated, where appropriate.

| **Species** | | **Goodenough** | **Fergusson** | **Normanby** |
| --- | --- | --- | --- | --- |
| Spotted Whistling-Duck *Dendrocygna guttata* | N | | Y | N |
| Radjah Shelduck *Radjah radjah radjah* | Y | | Y | N |
| Orange-footed Megapode *Megapodius reinwardt* *macgillvrayi* | Y | | Y | Y |
| **Blue-breasted Quail *Synoicus chinensis*** | N | | Y | N |
| Metallic Pigeon *Columba vitiensis* | Y | | Y | N |
| Amboyna Cuckoo-Dove *Macropygia amboinenesis* | Y | | Y | Y |
| Black-billed Cuckoo-dove Macropygia nigrirostris | Y | | Y | N |
| Great Cuckoo-Dove *Reinwardtoena reinwardtii* | Y | | Y | N |
| Pacific Emerald Dove *Chalcophaps longirostris* | Y | | Y | Y |
| Stephan’s Dove *Chalcophaps stephani* | Y | | Y | Y |
| White-bibbed Ground Dove *Pampusana jobiensis jobiensis* | Y | | Y | N |
| Nicobar Pigeon *Caloenas nicobarica* | Y | | Y | Y |
| Pheasant Pigeon *Otidiphaps nobilis insularis* | N | | Y | N |
| Pink-spotted Fruit-Dove *Ptilinopus perlatus zonorus* | Y | | Y | Y |
| Orange-fronted Fruit-Dove *Ptilinopus aurantiifrons* | N | | Y | Y |
| Wompoo fruit-dove *Ptilinopus magnificus* | N | | N | Y |
| Superb Fruit-Dove *Ptilinopus superbus* | Y | | Y | Y |
| White-breasted/Mountain Fruit-Dove *Ptilinopus rivoli* *bellus* | Y | | Y | N |
| Claret-breasted Fruit-Dove *Ptilinopus viridus vicinis* | Y | | Y | Y |
| Island Imperial-Pigeon *Ducula pistrinaria postrema* | Y | | Y | Y |
| Pinon’s Imperial-Pigeon *Ducula pinon salvadorii* | Y | | Y | Y |
| Zoe’s Imperial-Pigeon *Ducula zoeae* | Y | | Y | Y |
| Torresian Imperial-Pigeon *Ducula spilorrhoa* | Y | | Y | Y |
| Papuan Mountain-Pigeon *Gymnophaps albertisii* | Y | | Y | N |
| Pheasant Coucal *Centropus phasianinus* | Y | | Y | Y |
| Dwarf Koel *Microdynamis parva* *parva* | Y | | Y | Y |
| Channel-billed Cuckoo *Scythrops novahollandiae* | Y | | Y | Y |
| Shining Bronze-Cuckoo *Chrysococcyx lucidus* | Y | | Y | Y |
| Little Bronze Cuckoo *Chrysococcyx minutillus* | Y | | Y | Y |
| Brush Cuckoo *Cacomantis variolosus* | Y | | Y | Y |
| Marbled Frogmouth *Podargus ocellatus intermedius* | Y | | Y | Y |
| Large-tailed nightjar *Caprimulgus macrurus* | Y | | Y | Y |
| Barred Owlet-nightjar *Aegotheles bennetti plumifer* | Y | | Y | Y |
| Glossy Swiftlet *Collocalia esculenta* | Y | | Y | Y |
| Mountain Swiftlet *Aerodramus hirundinaceu*s | Y | | N | N |
| Uniform Swiftlet *Collocalia vanikorensis* | Y | | Y | Y |
| Beach Thick-knee *Esacus magnirostris* | Y | | Y | Y |
| Pied Stilt *Himantopus leucocephalus* | N | | Y | N |
| Little Ringed Plover *Charadrius dubius dubius* | Y | | N | N |
| Comb-crested Jacana *Irediparra gallinacea novaehollandiae* | Y | | Y | N |
| **Whimbrel *Numenius phaeopus*** | Y | | Y | N |
| Common Sandpiper *Actitis hypoleucos* | N | | Y | N |
| Lesser Frigatebird *Fregata ariel* | Y | | Y | Y |
| Oriental Darter *Anhinga melanogaster* | N | | Y | N |
| Little Black Cormorant *Phalacrocorax sulcirostris* | N | | Y | N |
| Little Pied Cormorant *Microcarbo melanoleucos* | N | | Y | N |
| Australian Pelican *Pelecanus conspicillatus* | Y | | N | N |
| Great-billed Heron *Ardea sumatrana* | N | | N | Y |
| Black Bittern *Ixobrychus flavicollis* | N | | N | Y |
| **Little Egret *Egretta garzetta*** | N | | Y | N |
| Pacific Reef-Heron *Egretta sacra* | Y | | Y | Y |
| Great Egret *Ardea alba* | N | | N | Y |
| **Plumed Egret *Ardea plumifera*** | N | | Y | N |
| Long-tailed Honey-buzzard *Henicopernis longicauda* | N | | Y | Y |
| Pacific Baza *Aviceda subcristata* | Y | | Y | Y |
| Gurney’s Eagle *Aquila gurneyi* | Y | | Y | Y |
| Variable Goshawk *Accipiter hiogaster pallidimas* | Y | | Y | Y |
| Gray-headed Goshawk *Accipiter poliocephalus* | N | | Y | Y |
| Osprey *Pandion haliaetus cristatus* | Y | | Y | Y |
| Black Kite *Milvus migrans* | Y | | Y | Y |
| Whistling Kite *Haliastur sphenurus* | Y | | N | N |
| Brahminy Kite *Haliastur indus* | Y | | Y | Y |
| White-bellied Sea-Eagle *Icthyophaga leucogaster* | Y | | Y | Y |
| Papuan Boobook *Ninox theomacha goldii* | Y | | Y | Y |
| Blyth's Hornbill *Aceros plicatus* | Y | | Y | Y |
| Common Kingfisher *Alcedo atthis* | Y | | Y | Y |
| Azure Kingfisher Ceyx azureus | N | | Y | Y |
| Little kingfisher *Ceyx pusillus* | Y | | Y | Y |
| Dwarf kingfisher Ceyx solitarius | N | | Y | Y |
| Forest Kingfisher *Todiramphus macleayii* | Y | | Y | Y |
| Sacred Kingfisher *Todiramphus sanctus* | Y | | Y | Y |
| Colonist Kingfisher *Todiramphus colonus* | N | | Y | Y |
| Beach kingfisher *Todiramphus saurophagus* | Y | | Y | Y |
| Yellow-billed Kingfisher *Syma torotoro ochracea* | Y | | Y | Y |
| Rainbow Bee-eater *Merops ornatus* | Y | | Y | Y |
| Dollarbird *Eurystomus orientalis waigiouensis* | Y | | Y | Y |
| **Australian Hobby Falco longipennis** | N | | Y | N |
| Peregrine Falcon *Falco peregrinus* | Y | | Y | Y |
| Australasian Swamphen *Porphyrio porphyrio* | Y | | N | N |
| White-browed Crake *Poliolimnas cinereus* | N | | Y | N |
| Pale-vented Bush-hen *Amaurornis moluccana* | N | | N | Y |
| **Red-necked Crake *Ralllina tricolor*** | N | | Y | Y |
| Sulphur-crested Cockatoo *Cacatua galerita* | Y | | Y | Y |
| Buff-faced Pygmy-Parrot *Micropsitta pusio* *harteri* | N | | Y | N |
| Eclectus Parrot *Eclectus roratus* | Y | | Y | Y |
| Red-cheeked Parrot *Geoffroyus geoffroyi* | Y | | Y | Y |
| Double-eyed Fig-Parrot *Cyclopsitta diopthalma virago* | Y | | Y | Y |
| Purple-bellied Lory *Lorius hypoinochrous* | Y | | Y | Y |
| Papuan Hanging-Parrot *Loriculus aurantiifrons* *meeki* | Y | | Y | N |
| South Papuan Pitta *Erythropitta macklotti finschii* | Y | | Y | N |
| Varied Honeyeater *Gavicalis versicolor* | Y | | Y | Y |
| Puff-backed Honeyeater *Meliphaga aruensis* | Y | | Y | Y |
| Brown-backed Honeyeater *Ramsayornis modestus* | Y | | Y | Y |
| Papuan black Myzomela *Myzomela nigrita* | Y | | Y | Y |
| Long-billed Myzomela *Myzomela longirostris* | Y | | Y | N |
| Tawny-breasted Honeyeater *Xanthotis flaviventer* *spilogaster* | Y | | Y | Y |
| Helmeted Friarbird *Philemon buceroides* | Y | | Y | Y |
| Large-billed Gerygone *Gerygone magnirostris proxima* | Y | | Y | Y |
| Black-faced Cuckooshrike *Coracina novaehollandiae* | Y | | Y | Y |
| Varied Triller *Lalage leucomela* *obscurior* | Y | | Y | Y |
| Common/Slender-billed Cicadabird *Edolisoma tenuirostris* | Y | | Y | Y |
| Gray-headed Cicadabird *Edolisoma schisitceps vittatum* | Y | | Y | Y |
| Variable Shrikethrush *Colluricincla fortis fortis* | Y | | Y | Y |
| Sclater's Whistler *Pachycephala soror* | Y | | N | N |
| Black-tailed Whistler *Pachycephala melanura* | N | | Y | Y |
| Gray Whistler *Pachycephala simplex* *brunnescens* | Y | | Y | Y |
| White-breasted Woodswallow *Artamus leucorynchus* | Y | | Y | Y |
| Hooded Butcherbird *Cracticus cassicus hercules* | Y | | Y | Y |
| Northern Fantail *Rhipidura rufiventris* | Y | | Y | Y |
| Louisiade Fantail *Rhipidura louisiadensis* | Y | | Y | Y |
| Willie-wagtail *Rhipidura leucophrys* | Y | | Y | Y |
| Spangled Drongo *Dicrurus bracteatus* | Y | | Y | Y |
| Trumpet Manucode *Phonygammus keraudrenii* | Y | | Y | Y |
| Curl-crested Manucode *Manucodia comrii* *comrii* | Y | | Y | Y |
| Goldie's Bird-of-Paradise *Paradisaea decora* | N | | Y | Y |
| Golden Monarch *Carterornis chrysomela* | Y | | Y | Y |
| Black-faced Monarch *Monarcha melanopsis* | Y | | Y | Y |
| Island Monarch *Monarcha cinerascens rosselianus* | Y | | Y | Y |
| Fan-tailed Monarch *Symposiachrus axillaris fallax* | Y | | N | N |
| Spectacled Monarch *Monarcha trivirgatus* | N | | N | Y |
| Spot-winged Monarch *Monarcha guttula* | Y | | Y | Y |
| Leaden Flycatcher *Myiagra rubecula* s*ciurorum* | N | | Y | Y |
| Satin Flycatcher *Myiagra cyanoleuca* | Y | | Y | Y |
| Shining Flycatcher *Myiagra alecto lucida* | Y | | Y | Y |
| Gray Crow *Corvus tristis* | Y | | Y | Y |
| Torresian Crow *Corvus orru* | Y | | Y | Y |
| Spectacled Longbill *Oedistoma ilolophus fergussonis* | Y | | Y | Y |
| Pygmy Longbill *Oedistoma pygmaeum meeki* | Y | | Y | Y |
| Torrent Flyrobin *Monachella muelleriana* | N | | Y | N |
| Golden-headed Cisticola *Cisticola exilis* | Y | | Y | Y |
| Pacific Swallow *Hirundo tahitica* | Y | | Y | Y |
| Mountain Leaf-Warbler *Phylloscopus trivirgatus* | Y | | Y | N |
| Black-crowned White-eye *Zosterops atrifrons* | Y | | Y | Y |
| Capped White-eye *Zosterops fuscicapilla crookshanki* | Y | | Y | N |
| Louisiade White-eye *Zosterpos griseotinctus* | N | | N | Y |
| Metallic Starling *Aplonis metallica* | Y | | Y | Y |
| **Singing Starling *Aplonis cantoroides*** | N | | Y | Y |
| Island Thrush *Turdus policephalus canescens* | Y | | N | N |
| Red-capped Flowerpecker *Dicaeum geelvinkianum violaceum* | Y | | Y | Y |
| Black Sunbird *Leptocoma aspasia christianae* | Y | | Y | Y |
| Sahul Sunbird *Cinnyris frenatus* | Y | | Y | Y |
| **Chestnut-breasted Munia *Lonchura castaneothorax ramsayi*** | Y | | Y | Y |
| Blue-faced Parrotfinch *Erythrura trichroa* | Y | | Y | N |
| Eurasian Tree Sparrow *Passer montanus* | N | | Y | N |
